# 2,4-Diacetylphloroglucinol Against *Candida albicans*: Biofilm Formation, Aspartyl Protease Production and Ultrastructure Changes

**DOI:** 10.1101/2022.04.26.489565

**Authors:** Artyom A. Stepanov, Alexey S. Vasilchenko

## Abstract

The dimorphic fungus *Candida albicans* is the one of the most important opportunistic pathogen of human. The use of fungicides against candida encounters sub-inhibitory effects, which are fungal stress response and undesirable for the host. In this work we investigated antifungal action of 2,4-diacetylphloroglucinol against *Candida albicans* ATCC 10231 with focus on they biofilm forming ability under the treatment.

It was found that 2,4-DAPG reduced ability of candida cells to form biofilm, but complete inhibition or irradiation did not achieve. Moreover, *Candida albicans* cells in the adherent state have been characterized by reduced susceptibility to 2,4-DAPG comparing to planktonic cells.

Investigation of mechanisms which could be explain the antibiofilm action of 2,4-DAPG has revealed the reduction in cell’s surface hydrophobicity and inhibition of yeast-to-hyphae transition. Biofilms formed under the sub-inhibitory concentrations of 2,4-DAPG were depleted in protein and carbohydrate content. Furthermore, we microscopically visualized the treated biofilms and have revealed numerous cannels localized on hyphae and associated with secretion of aspartyl proteases (Sap).

We assumed what excretion of Sap is triggered by reactive oxygen generated upon affecting of mitochondrion respiration by 2,4-DAPG. Introducing of antioxidant Trolox simultaneously with 2,4-DAPG lead to reducing of Sap production. Thus, production of aspartyl proteases is a one of undesirable side effect of *Candida albicans* treatment, but using 2,4-DAPG in combination with antioxidants is the solution to overcome it.

## Introduction

2,4-diacetylphloroglucinol (2,4-DAPG) is a small aromatic compound which produced as a second metabolite by *Pseudomonas* spp. (Keel et al. 1996). 2,4-DAPG is not a potent antibiotic comparable to amphotericin B, for example. However, it is capable of exhibiting activity over a wide range of suppressed organisms (Kankariya et al. 2019; Almario et al. 2017). The prevalence of genes involved in the production of 2,4-DAPG and other phloroglucinol derivatives is very high among various species of *Pseudomonas* genus (Troppens et al. 2013). In this regard, it can be concluded that 2,4-DAPG is an important metabolite, the role of which is not associated exclusively with killing effect.

Direct antimicrobial effect of 2,4-DAPG is realize towards planktonic bacterial cells (Troppens et al. 2013; Kamei et al. 2003; Couillerot et al. 2011) and its biofilms (Julian et al. 2020) due to hydrophobic interactions with biological membranes and its perturbation (Gong et al. 2016). While indirect antimicrobial effect of 2,4-DAPG could be associated with its activity as a chemical messenger. For example, it is known the ability of 2,4-DAPG to interfere with the expression of genes involved in biofilm formation of *Bacillus subtilis* (Powers et al. 2015) and inhibits production of quorum sensing autoinducer in *Pectobacterium carotovorum* population (Julian et al. 2020).

The question was arising regarding the action of 2,4-DAPG against fungi. It is known that 2,4-DAPG affects the energy production in yeast cells through impairing mitochondrial function by depolarization its membrane (Troppens et al., 2013; Kwak et al. 2011). Also, it is known disorganization of hyphal tips of *P. ultimum* with fungicidal concentration of 2,4-DAPG, and inhibition hyphal growth at sub-inhibitory (de Souza et al., 2003). Production of fusaric acid by several *Fusarium oxysporum* have been regulated with sub-inhibitory concentrations of 2,4-DAPG (Mayer et al. 2013).

*C. albicans* is an opportunistic human pathogen that frequently cause superficial infections, but in some cases may cause life-threatening systemic infections (Cavalheiro et al. 2018). Virulence of *C. albicans* associated with ability to biofilm formation. Biofilms of *C. albicans* are complex structures composed of yeast cells, pseudohyphae and hyphae surrounded by exopolymeric matrix (Cavalheiro et al. 2018). Biofilm form allows *C. albicans* to increase resistance to antifungals and remain hidden from host immune system. Most of the manifestations of candidiasis are associated with the formation of biofilms on medical devices like catheters, which represent an escalating problem in healthcare (Kojic et al. 2004).

Here we challenged the antifungal activity of 2,4-DAPG using the model yeast pathogen *Candida albicans* ATCC 10231. The 2,4-DAPG action was evaluated towards forming and matured biofilms with subsequent analysis of underling reasons. We focused on differences between intact and treated candida biofilms. Furthermore, we established the important morphological changes of candida hyphae upon 2,4-DAPG action, which did not have been described earlier. Revealed new hyphal structures could be associated with protease excretion and is a manifestation of undesirable sub-inhibitory effect of 2,4-DAPG. However, this side effect is possible to be eliminated.

## Materials and methods

### Strain and Media

*Candida albicans* ATCC 10231 was used as a model fungal microorganism. Before assays, *C. albicans* was cultured on YPD agar (1% yeast extract, 2% peptone, 2% dextrose, 1,5% agar) at 37°C for 24 h. Prior to each assay, a colony of *C. albicans* was transferred into YPD broth and incubated at 30°C overnight with the agitation of 100 rpm.

Commercially available preparation of 2,4-DAPG was used in this study (Abcam, UK).

### Determination of minimal inhibitory concentration

Determination of minimal inhibitory concentration (MIC) was performed following the Clinical and Laboratory Standards Institute guidelines for broth dilution antifungal susceptibility testing of yeasts (de Sousa et al. 2020; CLSI M27 document). The MIC was determined as the lowest concentration of antibiotic for which no visible fungal growth could be observed after 24 h of incubation.

### Biofilm inhibition and biofilm destruction assays

To investigate the biofilm inhibitory effect of 2,4-DAPG, *C. albicans* biofilms were produced on pre-sterilized, polystyrene, flat-bottom 96-well microtiter plates according to Khan et al. (2014) with some modifications.

Following the adhesion phase (i.e., 10^7^ cells incubated in 100 μl RPMI-1640 w/ L-glutamine and phenol red (Sigma-Aldrich, Germany) for 90 min at 75 rpm at 37°C), the cell suspensions were aspirated and each well was washed twice with 150 μl of PBS to remove loosely adherent cells. RPMI-1640 medium containing 2,4-DAPG was transferred into each of the washed wells up to final volume 100 μl. Solution of DMSO (5% vol/vol) in RPMI-1640 medium was taken as control sample.

Plates were incubated at 37°C in a shaker at 75 rpm for 24 and 48 hours, the culture medium was replenished daily. The metabolic activity of *C. albicans* biofilms was determined quantitatively using a 2,3,5-triphenyl-tetrazolium chloride (TTC) reduction assay (Pemmaraju et al. 2016). The total biomass of biofilms was visualized by crystal violet assay (Abirami et al. 2020).

### The effect of 2,4-DAPG on pre-formed biofilms

To investigate the effect of 2,4-DAPG on pre-formed biofilms, *C. albicans* biofilms were prepared for 24 h at 37°C in a shaker at 75 rpm as described above. Then, the wells were washed twice with PBS and fresh RPMI-1640 medium (100 μL) containing different concentrations of 2,4-DAPG was added and the plate was further incubated at 37°C in a shaker at 75 rpm up to 48 h. Solution of DMSO (5% vol/vol) in RPMI-1640 medium was taken as control sample.

The metabolic activity of the *C. albicans* biofilms and the total biomass of biofilms was determined quantitatively as described above.

### The effect of 2,4-DAPG on germ tube formation

The germ tube inhibition assay was performed in a 96-well microplate using *C. albicans* ATCC 10231 strain according to Zuzarte et al. (2011) with some modifications.

N-acetyl-D-glucosamine medium was prepared as follow: 0.5% N-acetyl-D-glucosamine (Sigma-Aldrich, USA), 0.5% peptone (Sigma-Aldrich, USA), and 0.3% KH2PO4 were dissolved in distilled water at a final volume of 100 mL, and filtered through PVDF-membrane with pores 0.22 μm (Millipore, US). The inoculum suspension was adjusted to 10^7^ CFU/mL in 0.01 M PBS, then have been added to 170 μL of N-acetyl-D-glucosamine media containing 2,4-DAPG. The wells without 2,4-DAPG were taken as negative controls. Microplates were incubated at 37°C at 150 rpm for 3 h.

After that, at least 100 yeast cells from each well were counted with a light microscope (Primo Star, Zeiss, Germany), and germ tube inhibition was calculated by following the formula (Okamoto et al. 1993):

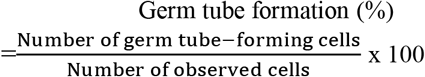

To evaluate hyphal growth of *C. albicans* ATCC 10231 strain on solid medium, 10 μL of inoculum suspension, adjusted to 10^6^ CFU/mL was spotted on Spider agar plates containing different concentrations (from 7,5 to 120 μg/mL) of 2,4-DAPG. The plates were incubated for 4 days at 37°C and the morphology of the fungal colony was observed using a stereomicroscope and photographed using a digital camera.

### Adhesion of *C. albicans* to polystyrene

Adhesion of *C. albicans* to polystyrene was performed on pre-sterilized, flat-bottom 96-well microtiter plates (Eppendorf, Germany) according to Vargas-Blanco et al. (2017) with some modifications. *C. albicans* ATCC 10231 was grown overnight in YPD broth, harvested, washed twice in PBS, and adjusted to 10^7^ CFU/mL and used for inoculation into a 96-well microtiter plate, containing RPMI-1640 medium and different concentrations (from 7,5 to 500 μg/mL) of 2,4-DAPG, for 90 min at 37°C.

RPMI-1640 medium with 5% DMSO was included in control wells. Following incubation, non-adherent *C. albicans* cells were removed by aspiration. The metabolic activity and total biomass of the adherent cells were determined quantitatively using a 2,3,5-triphenyl-tetrazolium chloride (TTC) reduction assay and crystal violet assay, respectively, as mentioned above.

### Cell surface hydrophobicity of *C. albicans*

Cell surface hydrophobicity (CSH) was assessed using the microbial adhesion assay to hydrocarbons (MATH) according to Salva-Dias et al. (2015) with some modifications. *C. albicans* was grown overnight at 37°C, were harvested and washed twice with PBS. Cell suspension displaying an optical density at 600 nm between 0,5 and 0,6 was prepared in PBS (A0); 2 mL of this yeast suspension was overlaid by 0,3 mL of the hydrophobic hydrocarbon, n-hexane. After vigorous vortexing (1 min), phases were allowed to separate (10 min) at 30°C and the optical density at 600 nm of the aqueous phase was measured (A1). The percentage of hydrophobicity was calculated by formula:

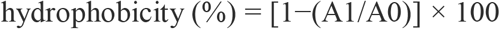

### Aspartyl protease activity of *C. albicans* planktonic cells and biofilms

Aspartyl protease activity was measured in two ways.

Production of SAP 1-3 subfamily was determined according to Jothi et al. (2021) with some modifications. Briefly, *C. albicans* planktonic cells and biofilms was grown in YPD medium supplemented with inhibitory or/and sub-inhibitory concentrations of 2,4-DAPG for 24 h. After incubation, the cell-free supernatant was collected through centrifugation at 8000 rpm for 10 min. Then, 0.1 mL of supernatant was mixed with 0.9 mL of citrate buffer (pH 3,5) consisting of 0.2 % of BSA. After incubation at 37 C for 15 min, the supernatant was precipitated with 15% TCA and centrifuged at 10000 rpm for 5 min. The concentration of the soluble products was measured at 280 nm.

Production of Sap 4-6 subfamily was measured according to Tamura et al. (2018) with some modifications. Briefly, 0.15 mL of cell-free supernatant was combined with 0.25 mL of 1% azocasein in Tris-HCl buffer (pH 5,5) and incubated at 37 °C for 1 h. Reaction was stopped with the addition of 0.4 mL 10% TCA. Subsequently, supernatant harvested by centrifugation (at 10,000×g for 10 min) was combined with an equal volume of 0.5 M NaOH, and incubated at 37 °C for 15 min. The concentration of the soluble products was measured at 440 nm.

### Lipase activity of *C. albicans*

Lipase activity was measured according to Ravindran et al. (2018) with some modifications. Briefly, *C. albicans* was grown in YPD medium supplemented with 1% Tween-20 and 2,4-DAPG at concentrations of 7.5, 15, 30, and 60 μg/mL for 24 h. After incubation, the cell-free supernatant was collected through centrifugation at 8000 rpm for 10 min. Then, 0.1 mL of supernatant was added to 0.9 mL of substrate mixture, which contained 1 volume of 0.3% p-nitrophenyl palmitate (pNPP) in isopropanol and 9 volumes of 0.2% Triton X-100 and 0.1% gummi arabicum in 25 mM Tris-HCl buffer (pH 8.0). Samples were incubated for 15 min at room temperature. After incubation, the reaction was terminated using 1M Na2CO3 and the absorbance was measured at 410 nm.

### Chemical analysis of biofilm matrix material

Extrapolymeric substances (EPS) from *C. albicans* biofilms extracted using formaldehyde/NaOH method (Liu and Fang 2002). Briefly, *C. albicans* biofilms were scrapped from microtiter wells, resuspended in PBS and vortexed at 300 rpm for 20 min. Biofilm suspension was first extracted with formaldehyde (36,5%) for 1 h and then with NaOH (1M, 4°C) for 3 h. All samples were centrifuged at 15000 rpm for 25 min and supernatants were filtered through 0.22 um membrane. The filtrates were used as EPS samples. Protein purification was performed with dialysis membrane (5 kDa) at 4°C for 24 h. Purified filtrate was frozen in liquid nitrogen and lyophilized. Freeze-dried pellet was resuspended in PBS at concentration 1 mg/mL. Carbohydrate content was measured with phenol-sulphuric method, protein content was measured with NanoPhotometer N120 (Implen, Germany).

### Microscopy investigations of *C. albicans* biofilms

#### Scanning electron microscopy

*C. albicans* biofilms were prepared on 8-well chamber slide (Eppendorf, Germany) according to Tsang et al. (2012) with some modifications. Briefly, slides were washed twice with PBS and dehydrated in a series of ethanol solutions (20% for 10 min, 50% for 10 min, 70% for 10 min and 96% for 15 min), and dried overnight in a lyophilizer (at 1,5 mbar) prior to sputter coating with gold. The surface topographies of the *C. albican*s biofilms were viewed with scanning electron microscope MIRA3 LMU (Tescan, Czech Republic).

#### Atomic force microscopy

The biofilms were formed as for SEM-investigation, but drying was performed on air during 12 hours. Atomic force microscopy was carried out using “Integra NT-MDT” (NT-MDT, Russia) microscope in tapping mode. Microscopy investigation was performed using the cantilever NSG01 (Tipsnano, Tallinn, Estonia) with spring constant ~5.1 N/m. Preliminarily, wide scanning of about 70 × 70 μm was conducted to produce a reference map of the sample surface so as to help localize the yeast cells. The scan size was then set with sampling of 512 by 512 points and scan rate of 0.5 Hz. Topographic, amplitude (MAG) and phase images were acquired simultaneously.

### Quantitative estimation of tyrosol production

*C. albicans* was grown in YPD at 30°C, 110 rpm for 24 h. Cells were harvested by centrifugation at 9000 g for 25 min. Supernatant were filtered through 0.22 um membrane. Tyrosol was isolated from culture supernatants by solid-phase extraction (SPE) according to Alem et al. (2006) with slight modifications. Tyrosol was quantified by reverse-phase high-pressure liquid chromatography (RP-HPLC).

Prior to extraction, 100 mL of cell free supernatant was acidified by 0.1 mol L^-1^ H_2_SO_4_ (prepared by adding 0.272 mL of 98% H_2_SO_4_ in 50 mL of distilled water). Extraction was performed with Strata C18 cartridge (Phenomenex, US).

To quantify the amount of tyrosol in the extracts, the analytical RP-HPLC was performed using Luna C-18 analytical column (4,6 × 250 mm, 5 μm, Phenomenex, US). Elution was performed using solvent B (80% acetonitrile in ultrapure water with 0.1% TFA) in a linear gradient according to the following scheme: 0–5 % for 5 min, 5-10 % for 5 min, 10 % for 5 min, 10-20% for 20 min, 20-70% for 5 min at a flow rate of 1.0 ml/min. Absorbance was detected at 225 nm.

OpenLab CDS ChemStation software was used for processing of the HPLC chromatograms. Calibration curve for tyrosol concentrations was plotted with 10, 20, 30, 50, and 100 μM of tyrosol which was purchased from Sigma-Aldrich (US).

### Statistical processing

The obtained results were statistically manipulated using Origin 2021 (OriginLab Corporation, Northampton, MA, USA) software.

The Shapiro–Wilk test was used to assess the normality of value distributions. In the presence of a normal distribution, the pair-sample Student’s t-Test was used.

Differences were considered significant at a p-value of 0.05. The data were presented as the mean and standard deviation (mean ±SD).

## 2 Results

### 2.1 Anti-biofilm properties of 2,4-DAPG

#### 2,4-DAPG inhibits formation of C. albicans biofilm

The effect of 2,4-DAPG on biofilms was evaluated at concentrations corresponding to the minimally inhibitory concentration (MIC) for planktonic cells (Fig. S1). It was found, that *Candida albicans* biofilms are more resistant to 2,4-DAPG than planktonic cells. 2,4-DAPG was unable to complete inhibit growth of biofilm at any used concentration, but was able to reduce its biomass and metabolic activity. 2,4-DAPG taken at 500 μg/ml halved biofilm biomass by the 24th hour (Fig. 1 a), while 125 μg/ml 2,4-DAPG decreased in biofilm biomass and metabolic activity for 50 % by the 48th hour (Fig. 1 b).

**Figure 1.**
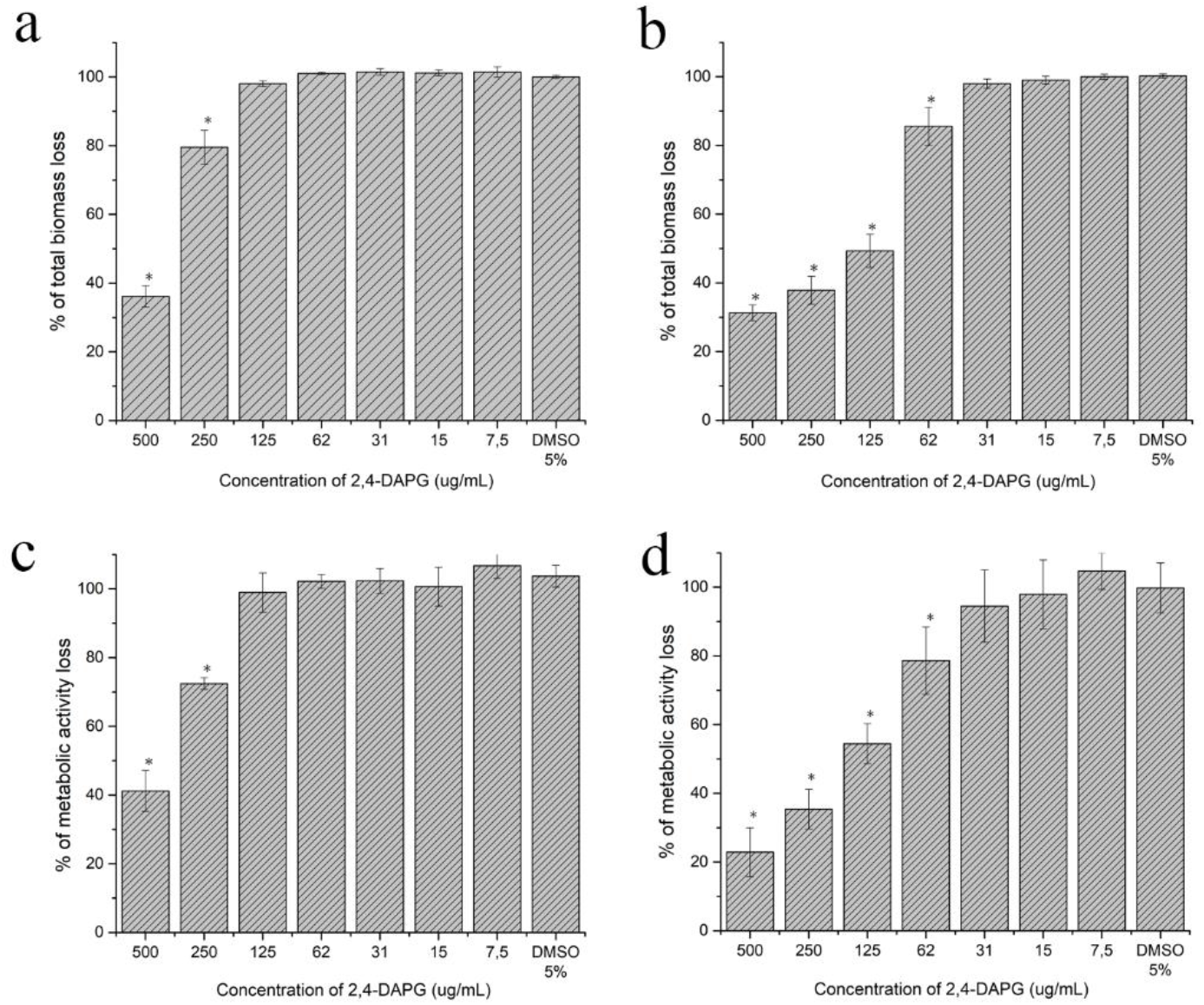
Influence of 2,4-DAPG on *C. albicans* ATCC 10231 biofilm development and biofilm metabolic activity. 2,4-DAPG action was estimated on biofilm biomass (a, b) and metabolic activity (c, b) in time-interval 24 hours (a, c) and 48 hours (b, d). Total biofilm biomass was quantified by crystal violet assay, while biofilm metabolic activity by TTC assay. Asterisk marks significance of differences between untreated (control) and each of 2,4-DAPG-treated samples (p <0.05, pair-sample Student’s t-Test).

The assessment of the metabolic activity of biofilm-embedded candida cells indicate that reducing in biofilm biomass is associated with decreasing in number of cells rather than amount of extracellular matrix (Fig. 1 c, d).

#### Effects of 2,4-DAPG on the preformed C. albicans biofilm

It was shown that 2,4-DAPG is also able to affect viability of pre-formed biofilms of *C. albicans.* 2,4-DAPG at concentrations 125 and 250 μg/mL reduces the metabolic activity of biofilm-embedded cells by 17.5 and 34.8%, respectively (Fig. 2 a). The more pronounced inhibition of metabolic activity (by 56%) was reached when mature biofilms were exposed to 500 μg/mL of 2,4-DAPG. However, eradication effect has not been revealed at all taken concentrations (Fig. 2 b).

**Figure 2.**
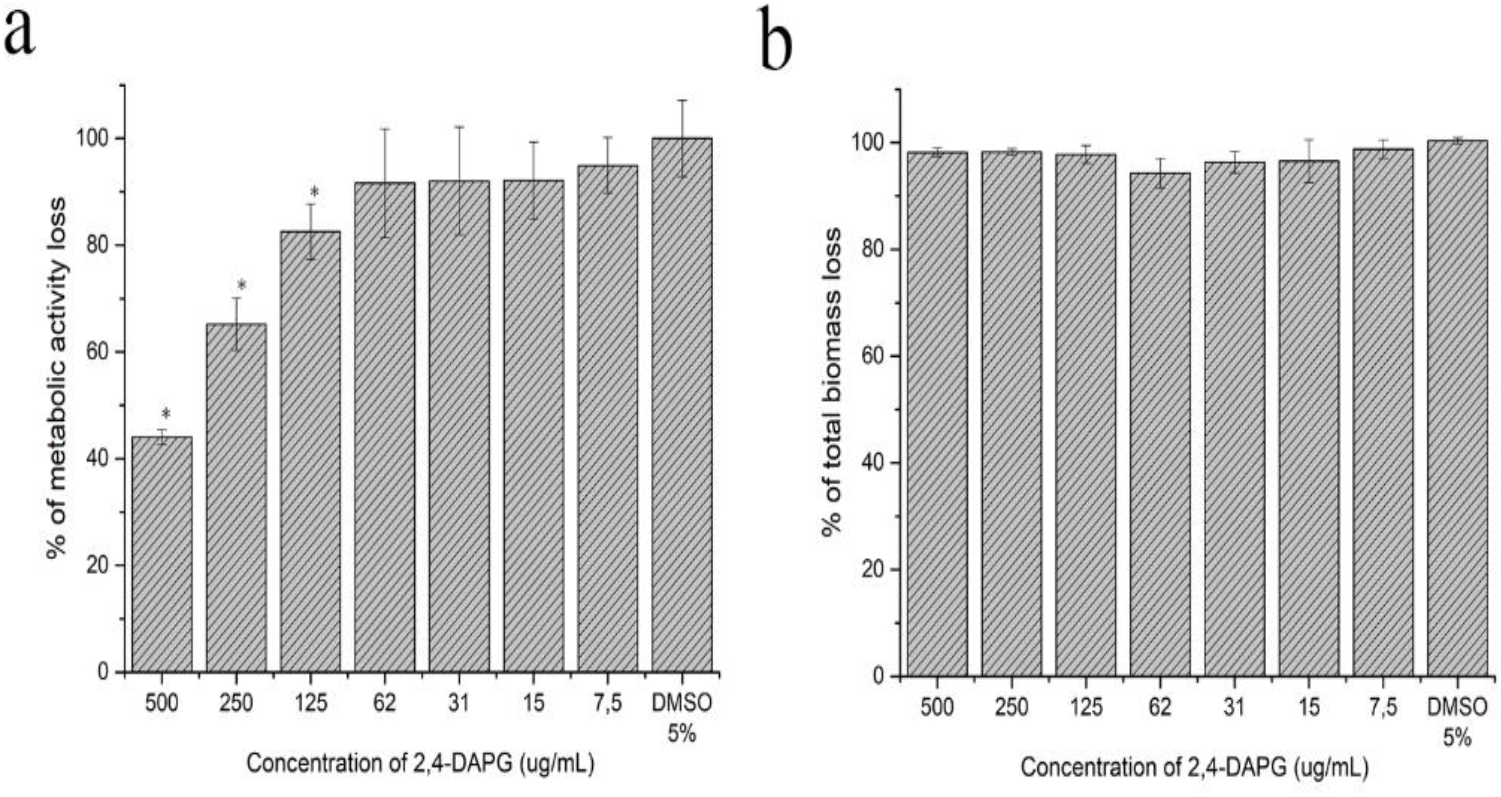
Influence of 2,4-DAPG on the structure of *C. albicans* ATCC 10231 mature biofilm and metabolic activity of biofilm-embedded cells. Metabolic activity was quantified by TTC assay (a). Total biomass of 48h mature biofilm was quantified by crystal violet assay (b). Asterisk marks significance of differences between untreated (control) and each of 2,4-DAPG-treated samples (p <0.05, pair-sample Student’s t-Test).

### 2.2. Microscopy investigations of *C. albicans* biofilms

Biofilms were formed on the surface of glass slides in the presence of various concentrations of 2,4-DAPG and studied using scanning electron microscopy (SEM). The biofilm morphology changed in two manner. The major changes were regarding to the thickness of biofilms and their structure. Thus, it was found that the maturing degree of biofilm is in relation with concentration used in co-incubation (Fig. S2). Concentrations 125-250 μg/ml decreased ability of *C. albicans* to form multilayer matured biofilm. Biofilms visualized with SEM as a monolayer of adherent cells with short hyphae which spatially distributed on a glass surface (Fig. 3c).

**Figure 3.**
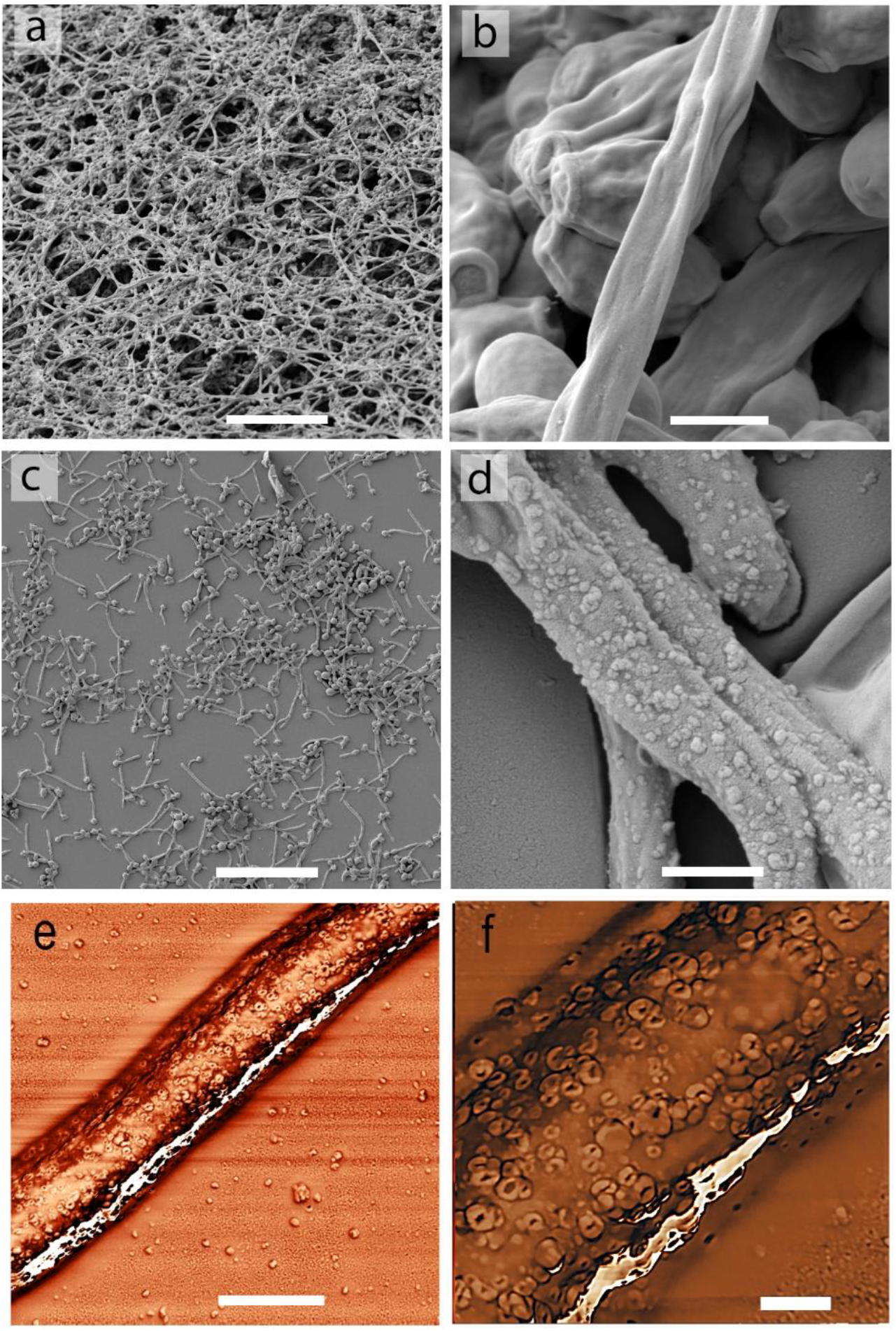
Microscopy investigations of *C. albicans* biofilm. a, b – SEM-images of the intact biofilm structure. c, d - SEM-images of the biofilm formed at the presence of 2,4-DAPG (250 μg/mL). e, a – AFM-images (phase shift mode) of hyphae treated with 2,4-DAPG (250 μg/mL) (e), and enlarged area of hyphae surface with channels (f). Scale bars - 50 μm (a, c); 2 μm (b, d, e); 200 nm (f).

A more detailed study of cells morphology revealed the tiny differences between normal and treated candida cells. It was shown, that hyphae but not budding cells contains numerous nanopores on their surface (Fig. 3 d). To exclude any SEM artifacts, we performed study of the biofilms with atomic force microscopy (AFM). It turned out that the surfaces of *C. albicans* hyphae grown in presence with 2,4-DAPG (125-250 μg/ml) become dotted (Fig. 3 e) with many channels with an average diameter of 172±42.6 nm (Fig. 3 f).

### 2.3. Chemical analysis of *C. albicans* biofilm’ matrix

To fully understand the effect of 2,4-DAPG treatment on biofilm structure, the analysis of extracellular polymeric substances (EPS) was performed. It was found reducing protein (Fig. 4a) and carbohydrate (Fig. 4b) content of biofilms, which were exposed to sub-inhibitory concentrations of 2,4-DAPG.

**Figure. 4.**
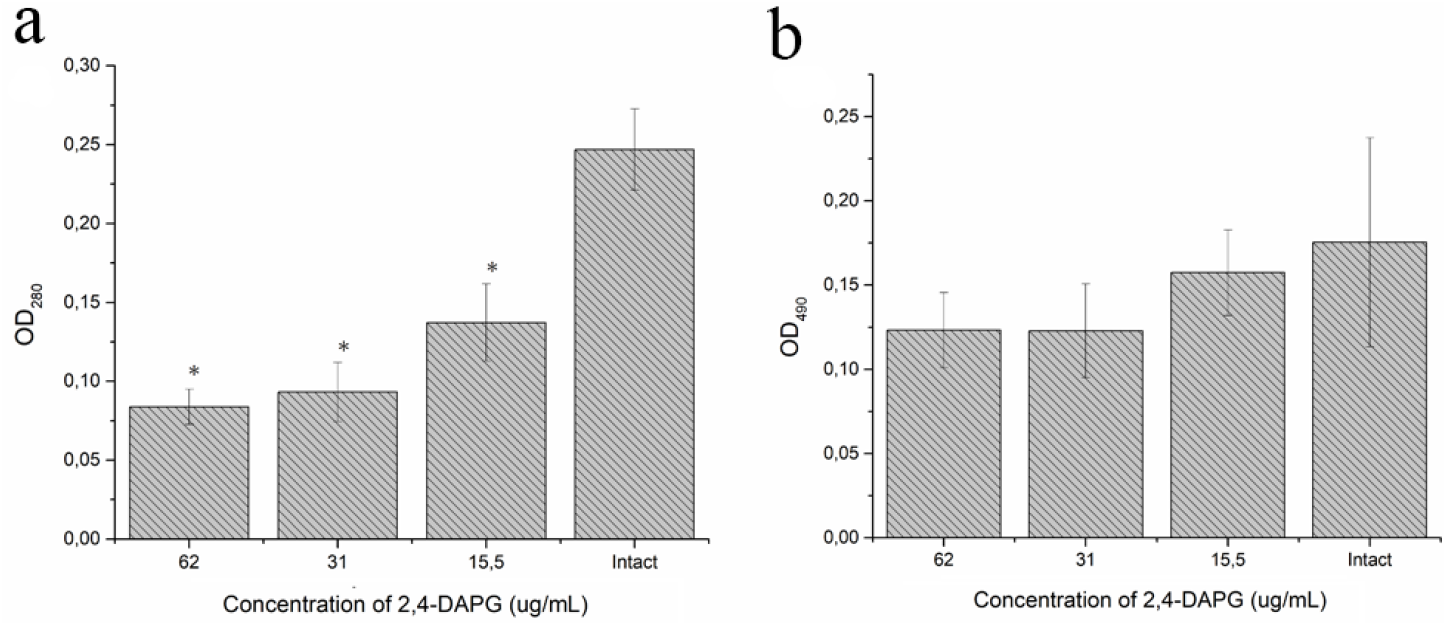
Protein and carbohydrate content in biofilm’s EPS of *C. albicans* ATCC 10231 exposed to sub-inhibitory concentrations of 2,4-DAPG. a – content of proteins, b – content of carbohydrates. Asterisk marks significance of differences between untreated (control) and each of 2,4-DAPG treated samples (p <0.05, pair-sample Student’s t-Test).

### 2.4. Investigation of biofilm inhibitory mechanisms of 2,4-DAPG

*Influence of 2,4-DAPG on the adhesive properties and surface hydrophobicity of C. albicans cells*

Assessing the ability of planktonic cells to adhere to the surface of a solid substrate, revealed reducing in the number of adherent cells by 47.5% and 17.4%, when 2,4-DAPG was applied for 500 and 250 μg/ml (Fig. 5a). Decreasing concentration more than 250 μg/ml had no effect on adhesion. At the same time, 60 and 30 μg/ml of 2,4-DAPG lowering the surface hydrophobicity of candida cell wall by 30.3 and 20%, respectively (Fig. 5b).

**Figure 5.**
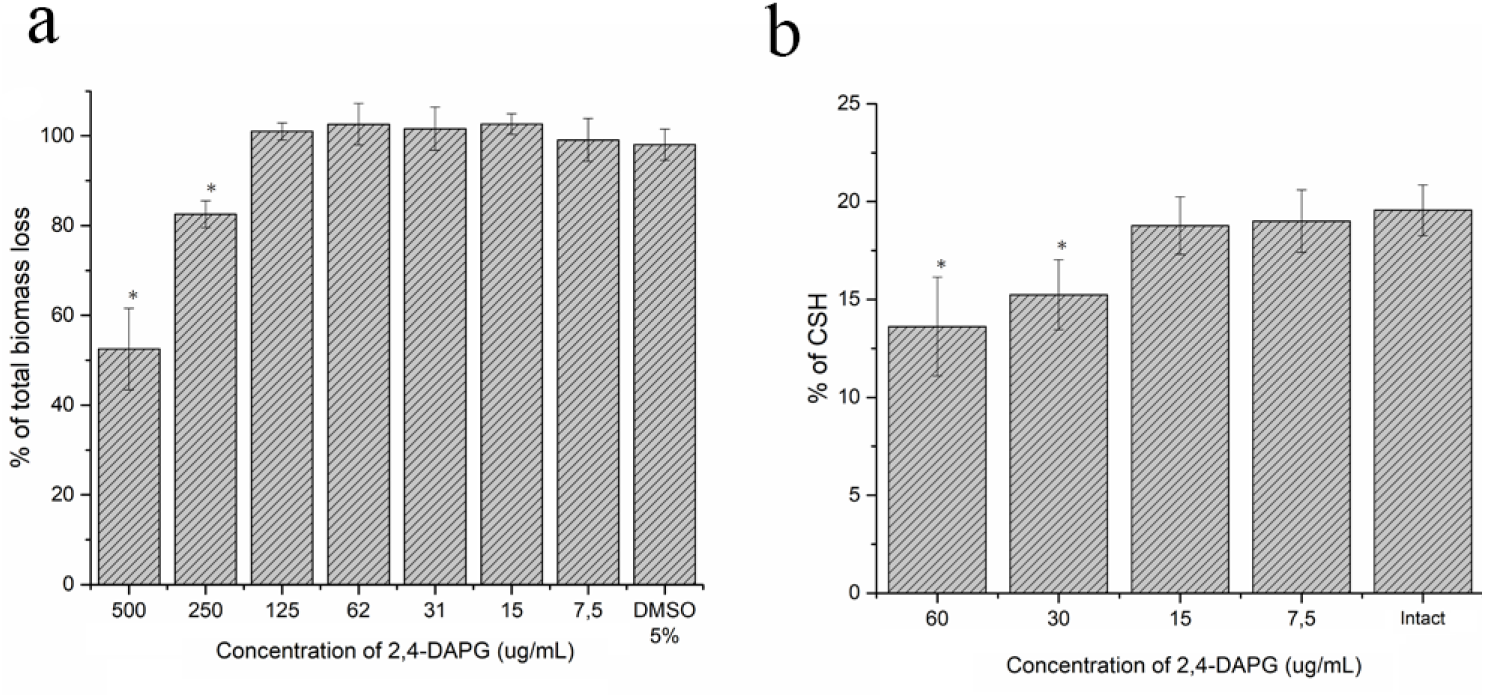
Adhesion to polystyrene and cellular surface hydrophobicity (CSH) of *C. albicans* ATCC 10231 cells. Total biomass quantification (crystal violet) assay was applied for estimation of the adhesion efficacy (a). Cell surface hydrophobicity of planktonic cells have been estimated by n-hexane assay (b). Asterisk marks significance of differences between untreated (control) and each of 2,4-DAPG-treated samples (p <0.05, pair-sample Student’s t-Test).

### 2.5. Effect of sub-inhibitory concentration of 2,4-DAPG on the aspartyl protease and lipase activities of planktonic and biofilm of *C. albicans*

It has been shown that sub-inhibitory concentrations of 2,4-DAPG do not affect lipase activity of *C. albicans* ATCC 10231 (Fig. S3).

The activity of aspartyl proteases 1-3 (Sap1-3) of *C. albicans* ATCC 10231 treated with sub-inhibitory concentration of 2,4-DAPG (60 μg/mL) was significantly higher comparing the untreated control. Overall aspartyl protease activity was increased by 15.7% (Fig. 6a).

**Figure 6.**
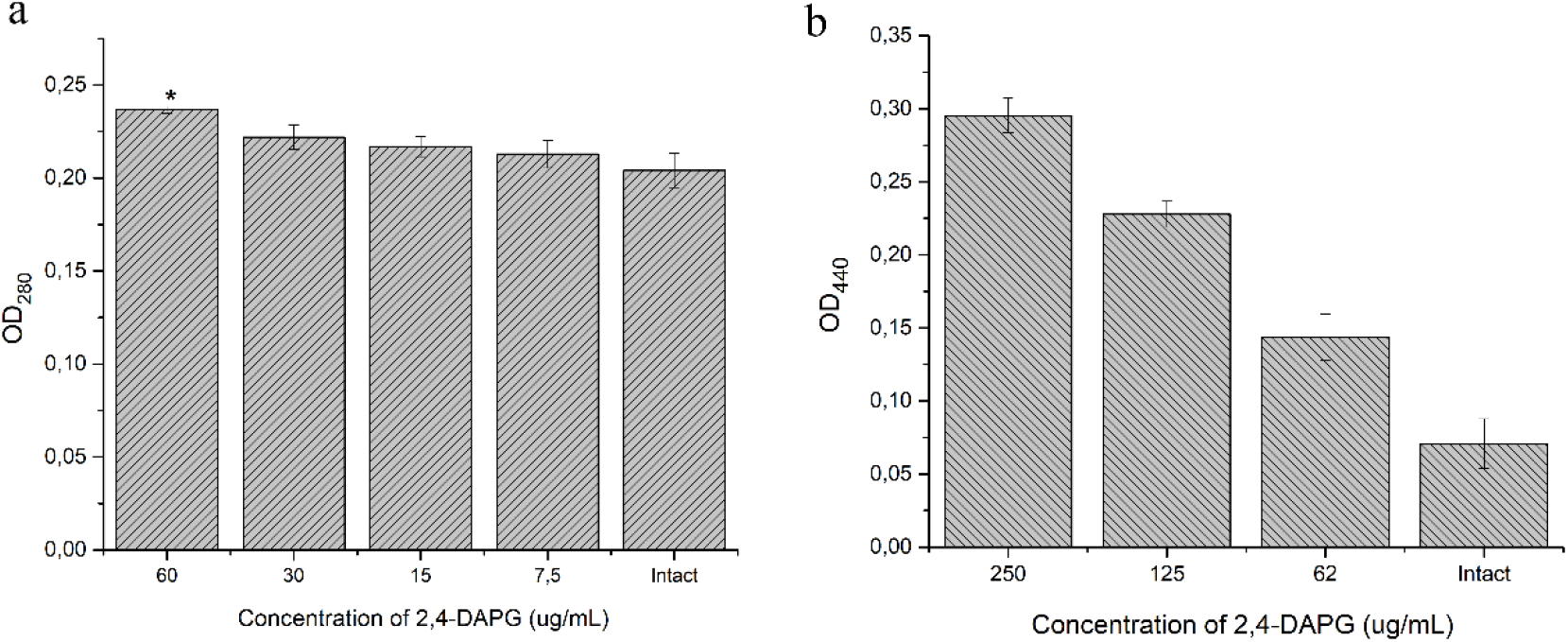
a) Protease (Sap1-3) activities of planktonic cells, b) and protease activity (Sap4-6) of biofilms *C. albicans* ATCC 10231 under the influence of different concentrations of 2,4-DAPG. Asterisk marks significance of differences between untreated (control) and each of 2,4-DAPG-treated samples (p <0.05, pair-sample Student’s t-Test).

The enhancement of aspartyl protease activity of *C. albicans* in presence of 2,4-DAPG forced us to check whether its secretion in biofilms. It was found no Sap1-3 activity was detected in either 2,4-DAPG-treated biofilms or control samples. However, it was shown that Sap 4-6 activity of *C. albicans* biofilms have been increased in 75.8, 68.2 and 54.4% when exposed to 2,4-DAPG in concentrations of 250, 125 and 62 μg/mL, respectively (Fig. 6b).

### 2.6. Effect of 2,4-DAPG on *C. albicans* dimorphism (germ tube formation/yeast-to-hyphae transition)

DAPG affects the morphogenesis of candida cells by suppressing the yeast-to-hyphae transition process. Hyphae formation in agar medium was completely inhibited when 2,4-DAPG was added for 30 μg/mL, and inhibited for 50 % at concentration 15 μg/ml (Fig. 7).

**Figure 7.**
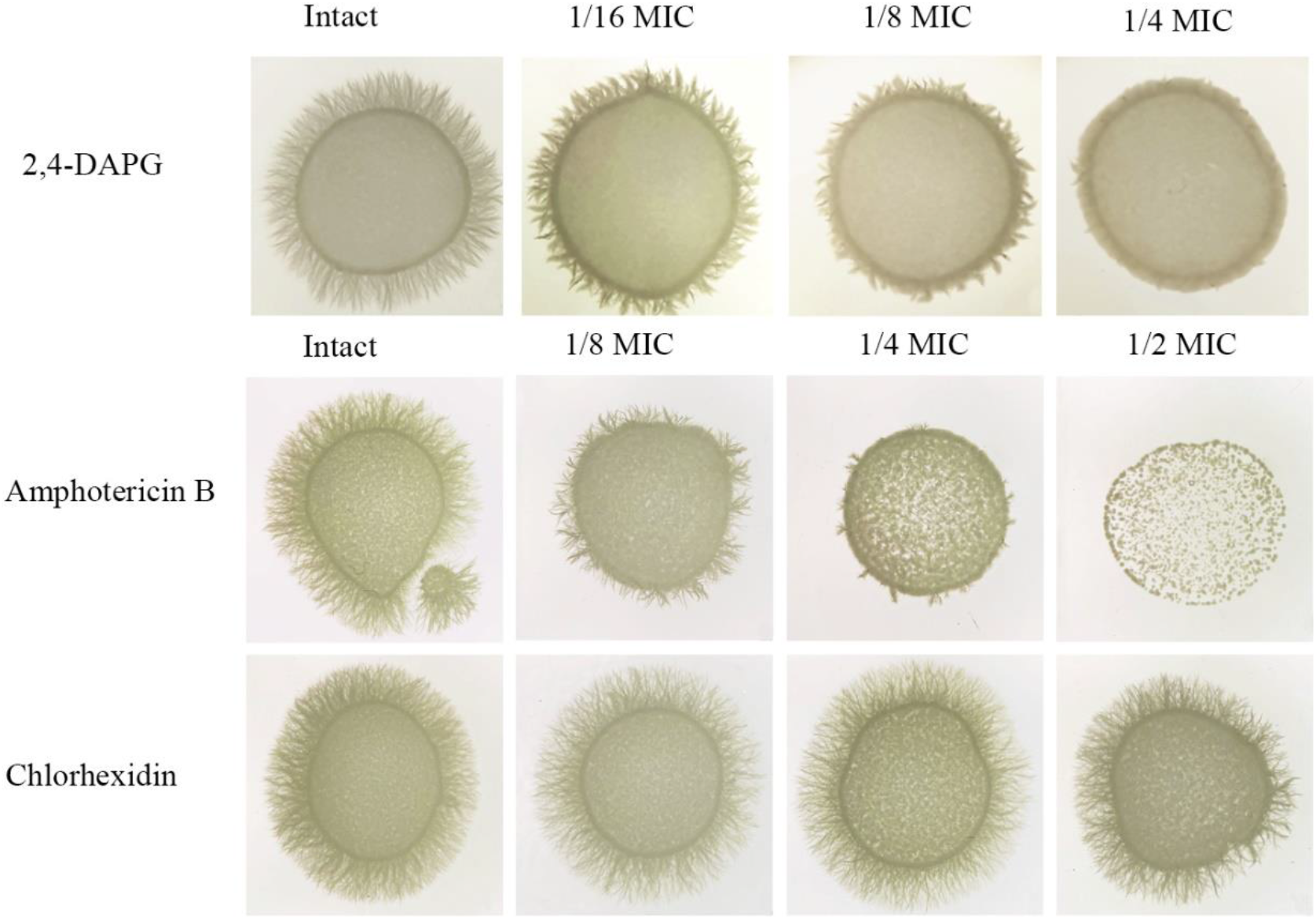
Yeast-to-hyphae transition of *C. albicans* ATCC 10231 on spider agar in the presence of sub-inhibitory concentrations various antifungal substances. 2,4-DAPG was added for 30 μg/mL (1/4 MIC); 15 μg/mL (1/8 MIC); 7.5 μg/mL (1/16 MIC); and intact control (0 μg/mL). Amphotericin B was added for 0.15 μg/mL (1/2 MIC), 0.075 μg/mL (1/4 MIC) and 0,037 μg/mL (1/8 MIC); chlorhexidine was added 0,0003% (1/2 MIC), 0,00015% (1/4 MIC) and 0,00007 (1/8 MIC).

Similar results were obtained with amphotericin B (Fig. 7). However, the observed suppression of *Candida albicans* cell morphogenesis is most likely not due to the toxic effect of 2,4-DAPG, since chlorhexidine taken at sub-inhibitory concentrations did not affects hyphal growth (Fig. 7).

Effect of 2,4-DAPG on morphogenesis of *C. albicans* was also evaluated in liquid medium. It was shown that addition of 2,4-DAPG in sub-inhibitory concentrations completely inhibit (30 μg/mL) or significantly reduce (15 μg/mL) germ tube formation (Fig. S4).

### 2.8. Influence of 2,4-DAPG on the production of quorum sensing autoinducers

Tyrosol is molecule that involve in quorum sensing controlled processes of Candida cells filamentation.

It was found that co-incubation of planktonic candida cells with sub-inhibitory concentrations of 2,4-DAPG leads to a dose-dependent decreased of tyrosol content in culture medium (Fig. 8 a). 2,4-DAPG in 15 and 30 μg/mL reduces tyrosol production for 62.8 and 76.8%, respectively, comparing to the untreated sample.

**Figure 8.**
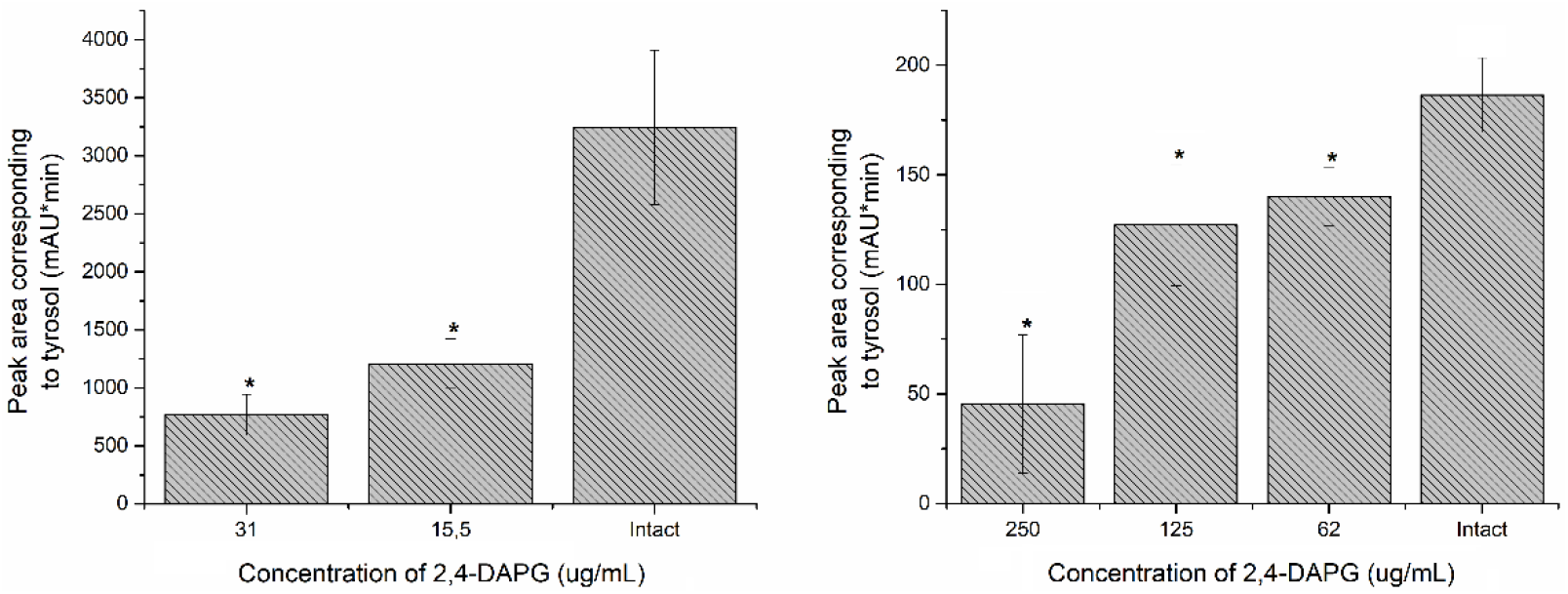
The amount of tyrosol in *C. albicans* culture media (a) and biofilm’ supernatant (b). Asterisk marks significance of differences between untreated (control) and each of 2,4-DAPG-treated samples (p <0.05, pair-sample Student’s t-Test).

Co-incubation of *C. albicans* biofilms with inhibitory and sub-inhibitory concentrations of 2,4-DAPG (62-250 μg/mL) also leads to decrease of tyrosol content in biofilm supernatant (Fig. 8b). 2,4-DAPG in 62, 125 и 250 μg/mL reduces tyrosol production for 25, 32, and 76%, respectively, comparing to the untreated sample.

In this regard, we tried to evaluate the effect of exogenous tyrosol on the ability of 2,4-DAPG to suppress hyphae formation. Using various combinations of tyrosol and 2,4-DAPG in the “checkerboard” assay, we failed to detect the restoring effect of exogenous tyrosol on *C. albicans* filamentation (data not shown).

### 2.9. Quantitative determination of 2,4-DAPG in *C. albicans* biofilm supernatant

To assess the ability of *C. albicans* biofilms to absorbance of 2,4-DAPG, a quantitative analysis of its residual amounts was carried out. It was found there was no significant difference in the residual amount of 2,4-DAPG between treatments with 250 and 125 μg/ml, while adding 60 μg/ml, the biofilms bound 93.6% (Fig. S5).

## Discussion

2,4-DAPG is a secondary metabolite that produces by wide variety of *Pseudomonas* species (Almario et al., 2017). In nature, the function of this molecule has not been reliably determined, but it is known that the production of 2,4-DAPG determines the properties of rhizospheric pseudomonads as agents of the biological control of phytopathogens (Weller et al. 2007; Brazelton et al. 2008).

It is also known that the antifungal effect of 2,4-DAPG on yeast cells is realized through disruption of mitochondrial membrane potential and uncoupling of oxidative phosphorylation (Troppens et al., 2013). The determined range of the minimum fungicidal concentration on planktonic yeast cells is 60-120 μg ml depending on the composition of the medium (Kwak et al. 2011),such MIC range agrees with our results.

The first thing we found is the resistance of *Candida albicans* ATCC 10231 cells to 2,4-DAPG action when they are in the state of adhesion to solid surface. Thus, we failed to achieve complete suppression of *Candida albicans* biofilm growth even when exposed to 500 μg/ml. Perhaps there is a significant restructuring of the metabolism of candida cells after their adhesion to a solid surface (Prażyńska and Gospodarek 2014). Most likely, this effect is not specific to 2,4-DAPG, since an increase in *Candida albicans* resistance when they exposed to amphotericin B and azoles are known (Prażyńska and Gospodarek 2014; Shuford et al. 2007). However, 2,4-DAPG significantly inhibits the growth and development of biofilms and is even capable of penetrating the matrix and suppress the metabolic activity of biofilm-embedded candida cells.

In the next stage, we analyzed the changes in the properties of planktonic cells in order to deep into biofilm inhibition mechanisms of 2,4-DAPG.

Cell surface hydrophobicity is an important biophysical property related with the microbial adhesion and biofilm formation (Danchik and Casadevall 2021; Borghi et al. 2011). It was found significant reducing in cellular surface hydrophobicity upon contact candida cells with 2,4-DAPG. It is known that

2,4-diacetylphloroglucinol is amphipathic molecule, which can not only enter through the lipid bilayer of cell membrane, but also changes hydrophobicity of the cell surface (Gong et al. 2016). Earlier, we found that local viscoelastic properties of bacterial surface are changed due to 2,4-DAPG treatment (Julian et al. 2020).

The second thing we found was the morphological changes in candida cells. Thus, during growth on a medium with 2,4-DAPG, the yeast-hyphal transition was disturbed. This process is directly related to the virulence of *C. albicans* and biofilm formation. The search for substances that is able to reduce virulence without a fungicidal effect is a perspective strategy for combating pathogens, since it promises to achieve a therapeutic effect without the resistance formation. There are numerous works dedicated to searching microbial or plant-derived molecules which inhibite hyphal formation and transform *C. albicans* to non-virulent phenotype (Atriwal et al. 2021).

In our study, 2,4-DAPG showed the ability to inhibit hyphal formation at sub-inhibitory concentrations when colony growth was unaffected. Similar effect was obtained with amphotericin B, for which the effect of inhibition of hyphae formation due to impaired ergosterol biosynthesis is well known (Kumar and Shukla, 2010; Nugent and Couchot 1986). On the contrary, sub-inhibitory concentrations of chlorhexidine taken as a control had no effect on hyphae formation.

Hyphal formation of *C. albicans* cells is complicated physiological process regulated through negative and positive expression of various genes. A significant role in hyphal formation is assigned to quorum sensing. *C. albicans* quorum sensing regulation is directed by small molecules tyrosol and farnesol, which biosynthesis started at log-phase and stationary phase, respectively (Alem et al. 2006; Hornby et al. 2001; Chen et al. 2004).

In our study, we found a significant decrease in tyrosol biosynthesis, but apparently this is do not determine the decrease in hyphal formation, since exogenous tyrosol did not restored germ tube formation. It’s known, tyrosol involved in acceleration of the morphological transition from yeast to hyphae, but not plays a major role (Alem et al. 2006). It is also need to note, we did not detect the production of farnesol, and did not revealed any effect on germ-tube formation in checkboard assay with 2,4-DAPG. However, *C. albicans* ATCC 10231 strain is known to produce farnesol at insignificant level (Riekhof and Nickerson 2007).

Thus, antibiofilm activity of 2,4-DAPG is realized through reducing candida cells adhesion, hydrophobicity and their morphological differentiation (hyphal formation) occurred at sub-inhibitory concentrations.

Analysis of the component composition of biofilms of 2,4-DAPG-treated cells revealed significant differences from intact biofilms. It’s known that matrix of *C. albicans* biofilms consists mainly of carbohydrates (30-41%) and proteins (3-5%), as well as a small amount of extracellular DNA (Paramonova et al. 2009). It was found that the content of proteins, but not carbohydrates, decreased in a dose-dependent manner in the 2,4-DAPG-treated biofilms. They may represent structural proteins (e.g. adhesins) or enzymes (e.g. proteases) (Flemming and Wingender 2010; Pinto et al. 2020; Nobile et al. 2008).

Studying biofilm morphology, we found some phenotypical changes that are presumably related to the defense response of *Candida albicans* to 2,4-DAPG. It was found that the action of 2,4-DAPG in concentrations of 125-250 μg/ml causes the formation of nanopores on the surface of hyphae. Similar formations, so-called “pimples”, were first described by Anderson et al. (1990) in *C. albicans* WO-1 isolated from immunosuppressed patient. Then, the enzyme-contained channels having a similar morphology to pimples were described at the *C. albicans* cell wall when grown on medium containing hydrocarbons (Zvonarev et al. 2017). The feature of 2,4-DAPG action is that nanopores were demonstrated exclusively on hyphae surfaces, but not on yeast cells. The function of nanopores is efflux of various biomolecules, including extracellular vesicles, which involved in virulence. Anderson et al. showed that *C. albicans* with pimples on cell surface, as well as vesicles hiding under them, secrete at least 10 times more aspartyl protease than phenotype without pimples (Anderson et al. 1990). Aspartyl proteases (Sap) constitute the main proteinase family directly involved in pathogenesis of *Candida* species (Ray et al. 1991).

We have checked the difference in aspartyl protease activity of *C. albicans* biofilm under the 2,4-DAPG-treatment. It was found that along with an increase in 2,4-DAPG concentration up to 250 μg/ml (by which nanopores are visualized) led to the increased activity of hyphae-associated Sap 4-6 activity. Thus, the formation of nanopores on the surface of hyphae upon exposure to high concentrations of 2,4-DAPG and the increase in aspartyl protease activity at the same concentrations is the interrelated phenomenon that indicates the pathogen’s response to the stress factors.

What is the trigger for increasing hyphae-associated Sap 4-6 activity under the influence of sub-inhibitory concentrations of 2,4-DAPG?

Primary mode of action of 2,4-DAPG lies in disruption of oxidative phosphorylation caused by changing mitochondrial membrane potential, whereby secondary mode of action includes production of reactive oxygen species induced by mitochondrial dysfunction leading to the peroxidation of intracellular molecules, along with acidification of cytoplasm caused by malfunction of vacuolar H^+^ ATPase (Fig. 9a) (Troppens et al. 2013; Kwak et al. 2011).

**Figure 9.**
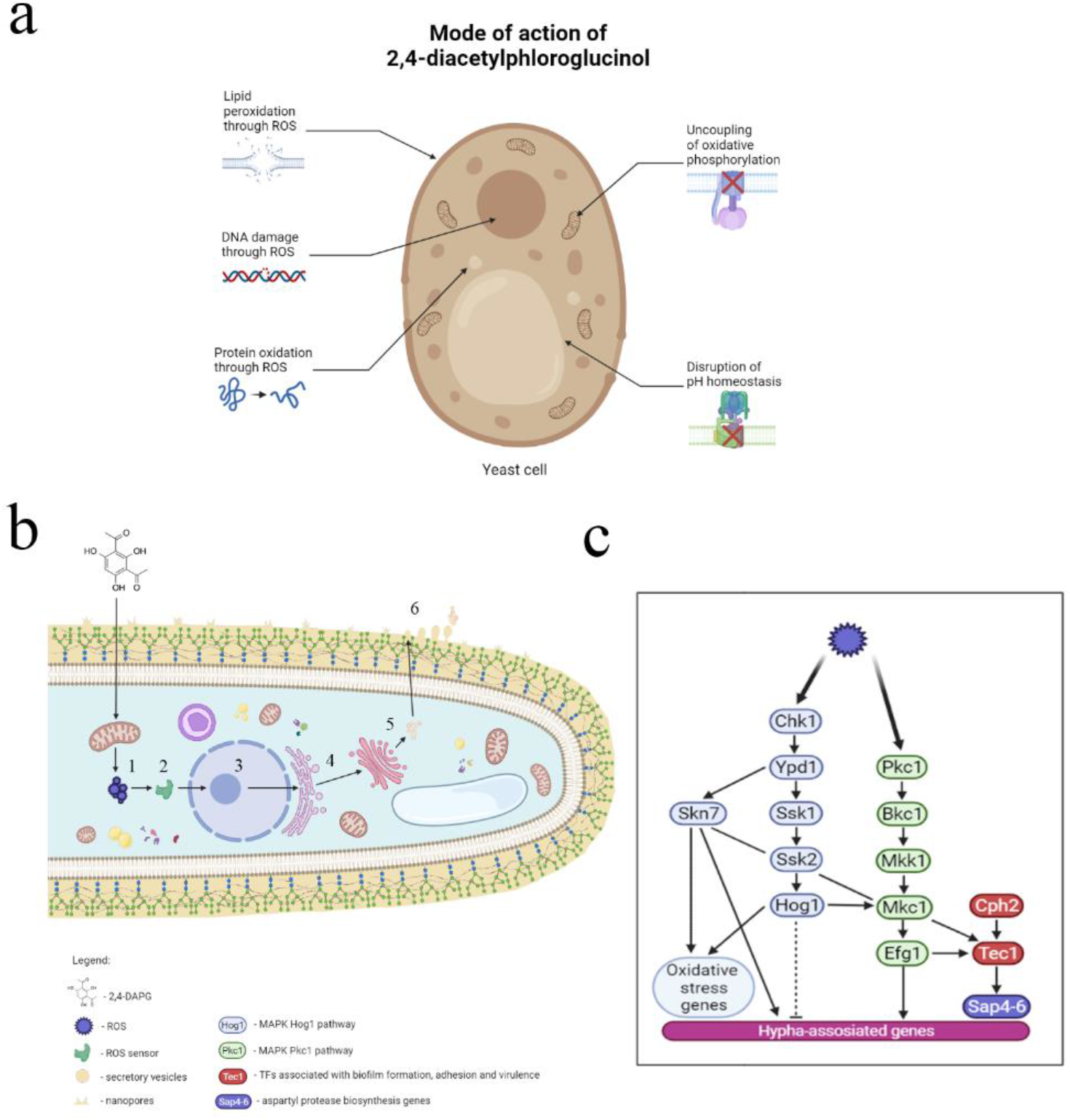
Mode of action of 2,4-DAPG on yeast cells, and its relationship with secretion of aspartyl proteases by *C. albicans*. 2,4-DAPG causes disruption of oxidative phosphorylation by changing mitochondrial membrane potential, increases production of reactive oxygen species leading to the peroxidation of intracellular molecules (a). Mode of action of 2,4-DAPG affects production of *Candida albicans* aspartyl proteases and realized through mitochondrial dysfunction. 2,4-DAPG affects mitochondria and generates ROS (1); ROS sensors (2) in cascade reactions activate expression of *sap 4-6* and hypha-associated genes (c) (3); the result of genes expression is launching of preproenzyme biosynthesis on EPS and modification it in proenzyme (4); proenzyme converts to its active form and packaged into vesicles by Golgi apparatus (5); the final stage is vesicles excretion throughout cell wall channels (6).

Oxidative stress is known to have direct and indirect effects on eukaryotic cells (Fig. 9 b). The direct action of reactive oxygen species is to oxidize intracellular molecules (e.g. proteins, DNA, lipids of plasma membrane), while the indirect action is aimed at regulating gene expression through various regulatory systems (Fig. 9 c) (Farrugia and Balzan 2012).

DAPG due to its amphipathic properties penetrates through cell wall of *C. albicans*, and causes dysfunction of mitochondria leading to production of reactive oxygen species. Further, ROS sensors become activated, what led to adaptation of the yeast to oxidative stress, and, to the activation of hypha-associated genes, including responsible for the synthesis of aspartyl proteases and secretion into extracellular environment (Basso et al. 2018).

Moreover, 2,4-DAPG-induced malfunction in vacuolar H^+^ATPase which led to pH changes in both vacuolar space and cytoplasm also can alters aspartyl protease activity (Troppens et al. 2013; Palmer et al., 2005).

If production of reactive oxygen species is a trigger for Sap’s production, then the administration of antioxidants should affect the proteolytic activity of candida cells. We used Trolox, water soluble analog of vitamin E which exhibit antioxidant activity.

It’s shown that addition of trolox simultaneously with 2,4-DAPG significantly reduces Sap4-6 activity of *C. albicans* biofilms without affecting their metabolic activity (Fig. 10). At the same time, co-incubation of *C. albicans* biofilms with 2,4-DAPG in the presence and absence of sub-inhibitory concentrations of pepstatin A, aspartyl protease inhibitor did not reveal significant differences between each other in Sap4-6 activity (data not shown).

**Figure 10.**
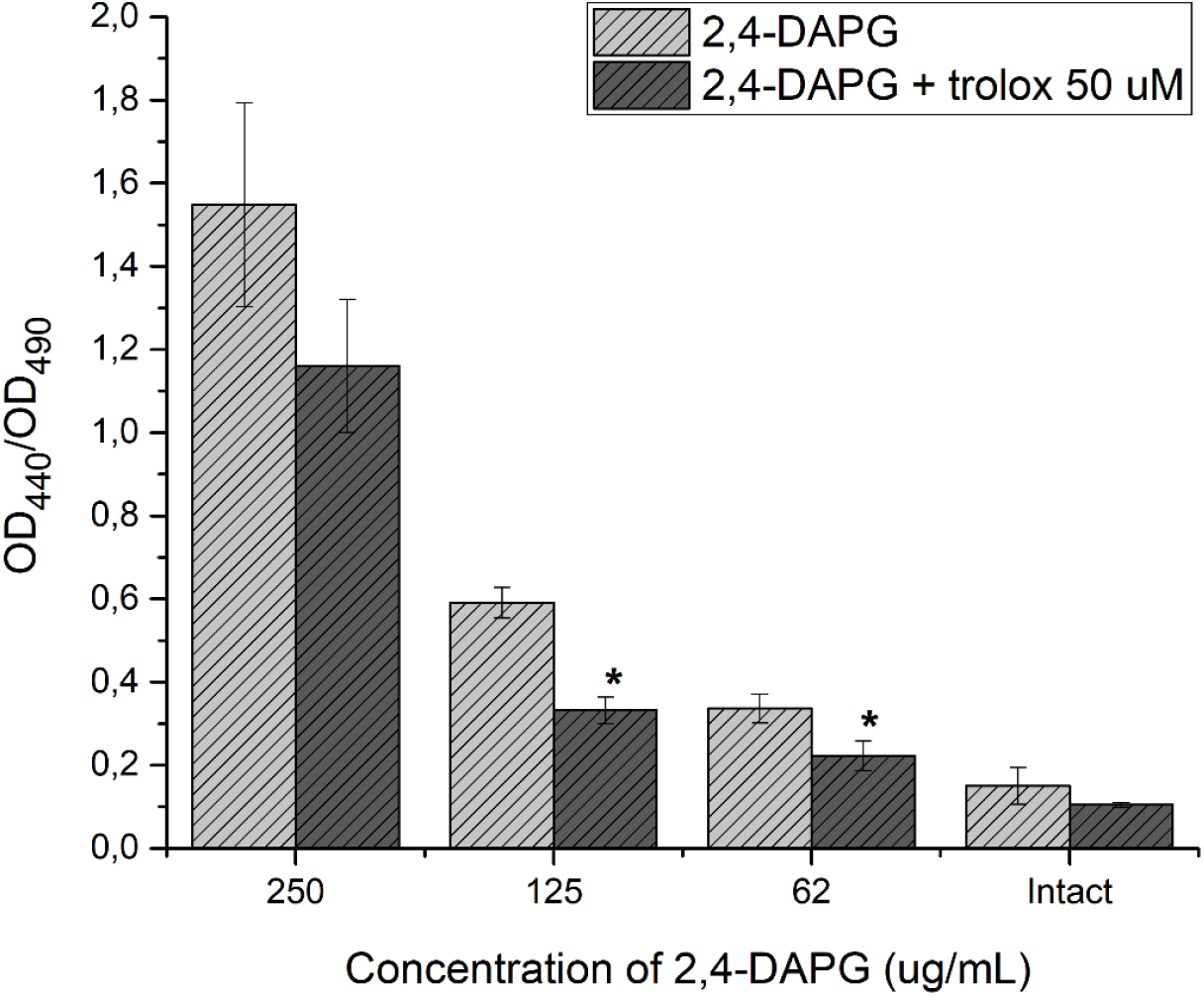
Aspartyl protease activity (Sap4-6) of *C. albicans* ATCC 10231 biofilms exposed to DAPG alone and in addition of trolox 50 uM.

Thus, enhanced aspartyl protease activity of *C. albicans* can be explained by the contribution of primary and secondary modes of action of 2,4-DAPG since increased production of ROS itself does not explain the reasons of this phenomenon.

