## Supplementary material for "2,4-Diacetylphloroglucinol Against *Candida albicans*: Biofilm Formation, Aspartyl Protease Production and Ultrastructure Changes": (Fig. S1)

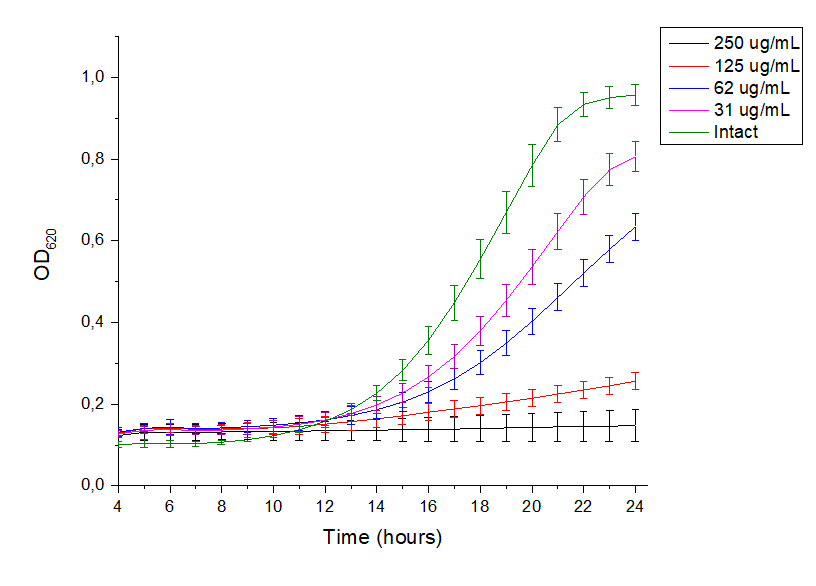
 Figure S1. Estimation of minimal inhibitory concentrations of 2,4-DAPG against planktonic cells of *C. albicans* ATCC 10231. Growth curves of *C. albicans* was built by recording the absorbance at 620 nm.

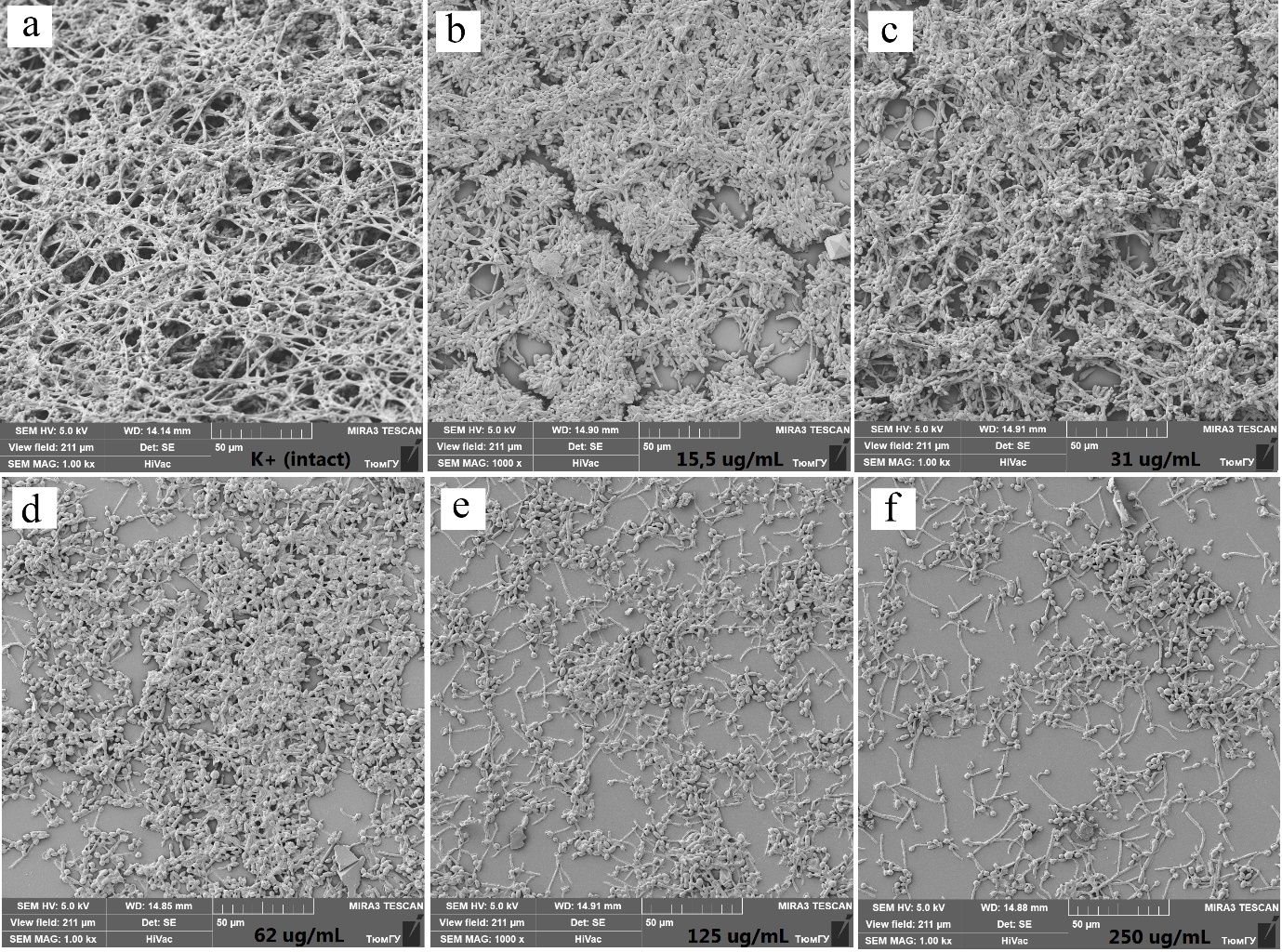

Figure S2. Scanning electron images of *C.albicans* ATCC 10231 biofilms, which were formed under the 2,4-DAPG influence: a- intact, b - 15.5 µg/mL, c- 31 µg/mL, d - 62 µg/mL, e - 125 µg/mL, f - 250 µg/mL.

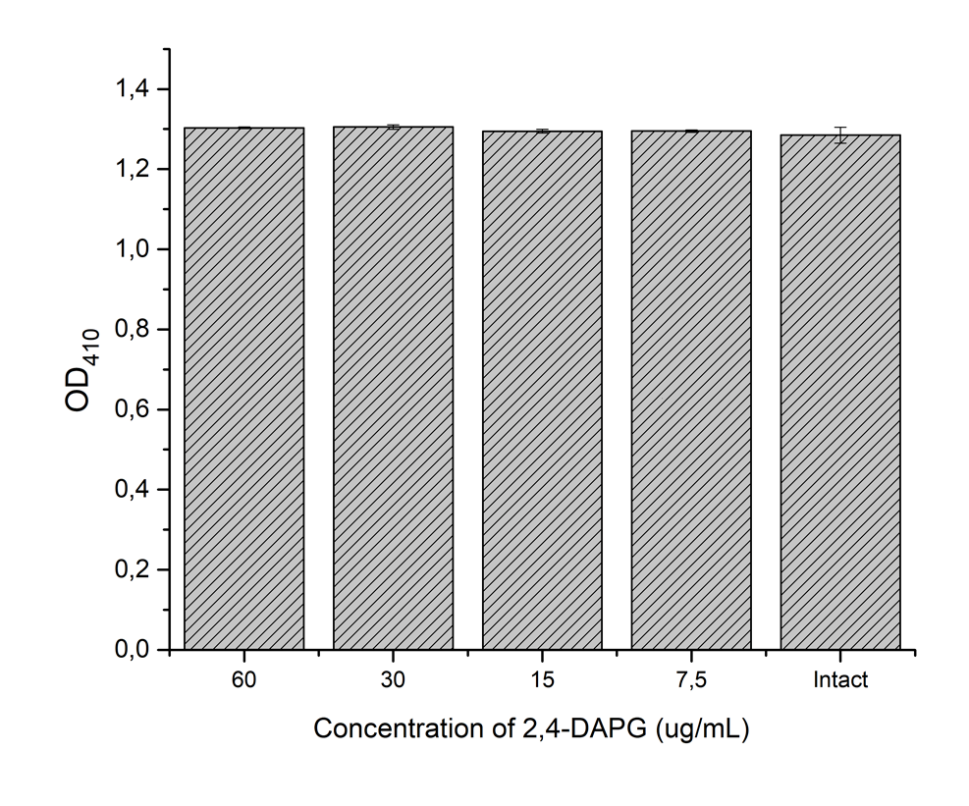

Figure S3. Lipase activity of planktonic cells of *C. albicans* ATCC 10231 exposed to different concentrations of 2,4-DAPG. Asterisk marks significance of differences between untreated (control) and each of 2,4-DAPG-treated samples (p <0.05, pair-sample Student’s t-Test).

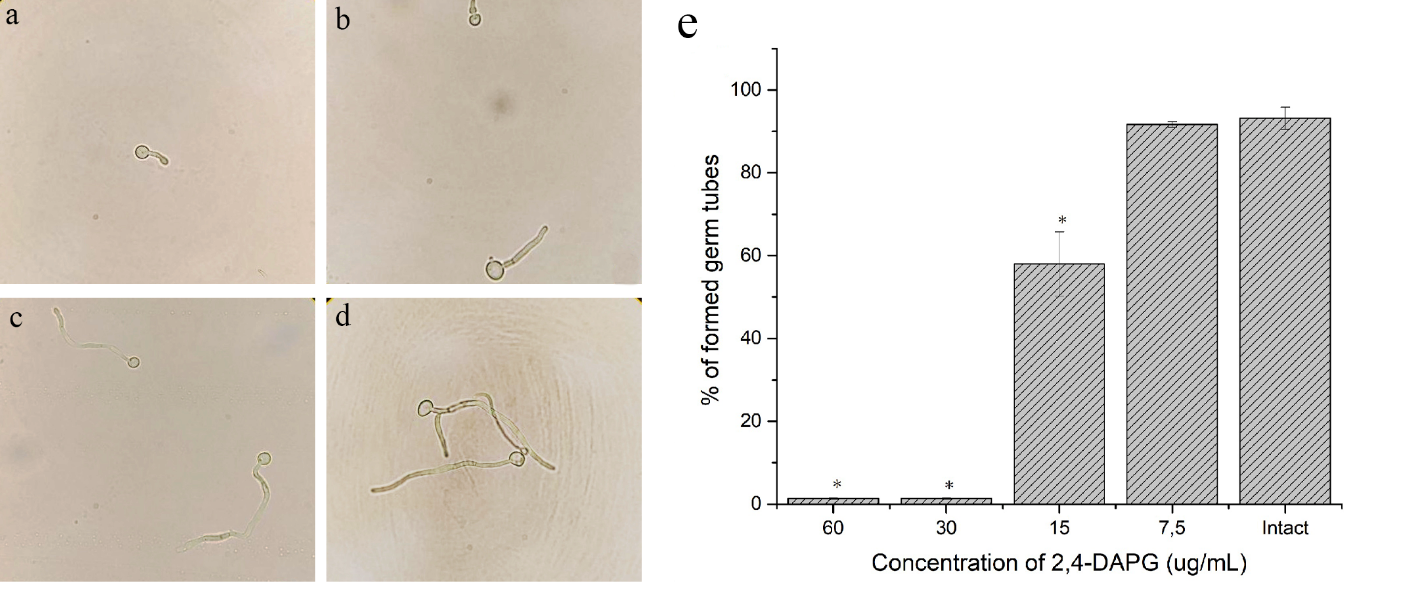

Figure S4. Inhibition of *C. albicans* ATCC 10231 filamentation.
Microscopic images showing morphology of *C. albicans* ATCC 10231 grown for 3 h under the 2,4-DAPG treatment with a) 61 µg/mL (½ MIC); b) 30 µg/mL ¼ MIC; c) 15 µg/mL ⅛ MIC; d) without 2,4-DAPG treatment. e) Percentage of formed germ tubes of *C. albicans* ATCC 10231 in the presence of different concentrations of 2,4-DAPG. Asterisk marks significance of differences between untreated (control) and each of 2,4-DAPG-treated samples (p <0.05, pair-sample Student’s t-Test).

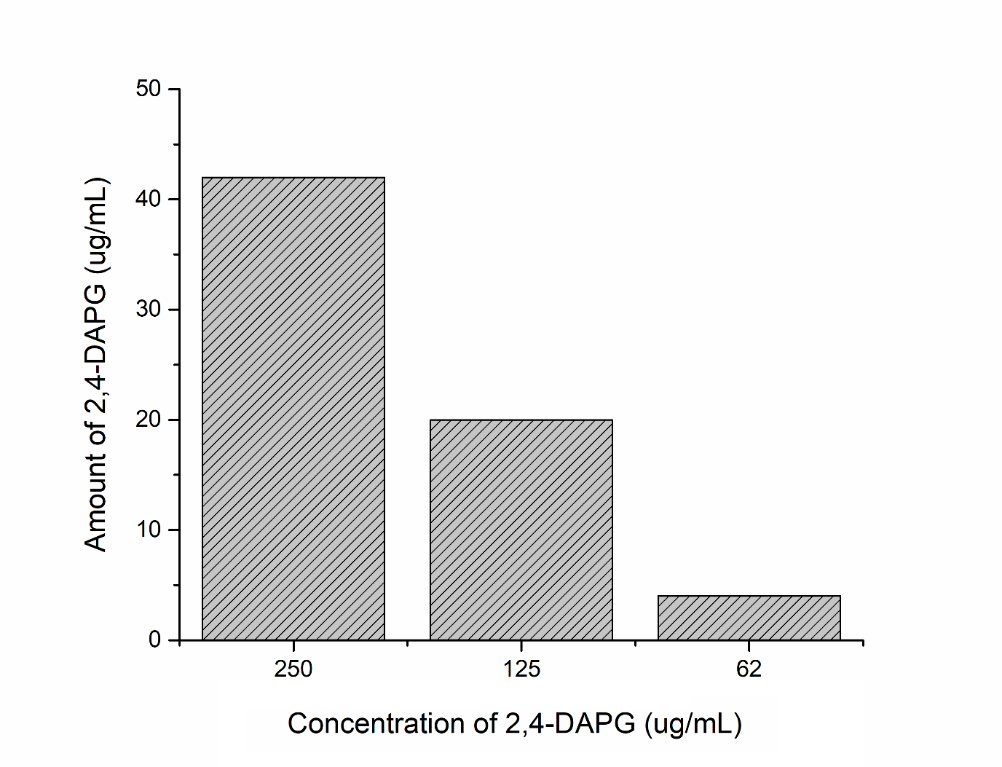

Figure S5. The ability of *Candida albicans* biofilms to absorb of 2,4-DAPG. The graph reflects the residual amount of 2,4-DAPG after co-incubation for 24 hours. The concentration was
